# α-Synuclein pathology differentially alters T-type calcium currents in vulnerable and resilient substantia nigra dopaminergic subpopulations

**DOI:** 10.64898/2026.07.10.737840

**Authors:** Megan L. Beaver, Pallavi Bommareddy, Natalie Z. McLean, Valerie J. Lewitus, Kathleen Maguire-Zeiss, Rebekah C. Evans

## Abstract

Aggregation of α-synuclein protein is a characteristic of Parkinson’s disease pathology that relates to the degeneration of vulnerable dopaminergic neurons and motor symptoms of the disease. However, α-synuclein pathology can contribute to neuronal dysfunction by disrupting several processes within the cell, including intracellular calcium balance, mitochondrial function, and synaptic function. Here, we use a preformed fibril (PFF) model of synucleinopathy to examine effects of striatal α-synuclein seeding on dopamine neurons of the substantia nigra *pars compacta* (SNc). The SNc is heterogeneous and contains dopaminergic neurons with differential vulnerability to Parkinson’s disease pathology. We found that intrastriatal injections of PFFs differentially affect these SNc neuron subtypes by increasing the excitability of resilient SNc neurons, while altering tonic firing patterns and T-type calcium currents in vulnerable SNc neurons. In addition, we performed comprehensive electrophysiological analyses and neural morphology reconstructions on SNc neurons from PFF and monomer injected mice. These findings provide insights to the selective vulnerability of SNc neuron subtypes and further our understanding of the role of α-synuclein in Parkinson’s disease progression and circuit dysfunction.

## Introduction

In Parkinson’s disease (PD), the progressive dysfunction and degeneration of dopaminergic neurons of the substantia nigra *pars compacta* (SNc) are associated with the formation of α-synuclein-comprised Lewy bodies^1^. Aggregated α-synuclein can spread from neuron to neuron through exo- and endocytic mechanisms^2,3^. In the cell, α-synuclein aggregates exert deleterious effects on multiple processes, causing mitochondrial dysfunction, impaired protein degradation, endoplasmic reticulum stress, membrane disruptions, and synaptic dysfunction^4–7^. Further, α-synuclein has been shown to inhibit dopamine transporter (DAT) and tyrosine hydroxylase (TH) activity, interfering with normal functioning of these cells^8–10^. Mutations in the gene expressing α-synuclein, SNCA, were the first to be linked to familial cases of PD^11^ and are also seen in spontaneous cases of PD. In familial cases of PD, it has been found that there is a dose-dependent correlation of α-synuclein load to phenotype severity^12^.

A long-standing question in PD research is why some dopaminergic neurons are more susceptible to neurodegeneration than others. It is well known that the ventral tegmental area (VTA) dopaminergic population is comparably resilient to neurodegeneration in PD compared to the SNc population^13–16^. However, within the SNc itself, there are vulnerable and resilient dopaminergic subpopulations^17–26^. Vulnerable SNc neurons are located in the ventral tier of the SNc and express Anxa1 and Aldh1a1, but lack the calcium buffering protein calbindin, while the resilient population is located in the dorsal tier and is positive for calbindin^17–26^. While these populations differ in their vulnerability to neurodegeneration, the relationship between these populations and α-synuclein remains unclear. Interestingly, one study examining human brain tissue from patients with dementia with Lewy bodies showed that Lewy body formation occurred exclusively in calbindin-negative neurons^27^, but there have been no comprehensive experiments in animal models comparing the effects of α-synuclein on the vulnerable and resilient populations of the SNc.

One important difference between the vulnerable and resilient SNc populations is the magnitude of T-type calcium current, which is larger in the vulnerable population^28^. Excessive calcium influx is an important aspect of selective vulnerability^29–31^, and intracellular calcium and α-synuclein have a reciprocally causal relationship^31–33^. High levels of calcium increase α-synuclein aggregation^34,35^ and α-synuclein aggregates disrupt calcium channel function and increase intracellular calcium^36–38^. Indeed, in human cell lines expressing α-synuclein, a transient increase of intracellular calcium increases α-synuclein aggregates^34,35^. Importantly, recent work has implicated T-type calcium channels for their role in PD^38,39^. T-type channels have 3 subtypes: Cav3.1, 3.2, and 3.3. Genetic mutations in T-type Ca^2+^-channel subtypes (Cav3.2, 3.3), but not L-type channels, were associated with PD status^39^. Over-expression of Cav3.2 or 3.3 increases cellular vulnerability in patient-derived cell cultures, while pharmacological inhibition or knockdown of any T-type calcium channel subtype is protective^38^. However, the influence of α-synuclein on T-type calcium currents in vulnerable and resilient SNc neurons is not understood. Here, we inject α-synuclein pre-formed fibrils (PFFs)^40–43^ to examine the effects of α-synuclein aggregation on vulnerable and resilient SNc dopaminergic subpopulations. Using a combination of *ex vivo* whole-cell patch-clamp electrophysiology, 3D morphological reconstruction, RNA fluorescence *in situ* hybridization, brain clearing, and immunohistochemistry techniques, we find that α-synuclein PFFs induce a time-dependent reduction in T-type calcium current selectively in vulnerable SNc neurons.

## Results

### Striatal injection of α-synuclein PFFs causes aggregation of phosphorylated α-synuclein within the SNc

To initiate α-synuclein pathology within the SNc, we injected wild-type mice with mouse α-synuclein PFFs or monomer controls in the dorsolateral striatum (see Methods for details). A total of 8 μg of protein was injected across four locations within the left hemisphere (Figure 1a). In this commonly used model^4,40,42,44–50^, α-synuclein travels up the axons to the SNc where phosphorylated α-synuclein (pα-syn)-immunoreactive aggregates and neurites can be identified in SNc neurons. Mice were euthanized at either 6- or 12-week timepoints post-injection for electrophysiological or RNAscope experiments (Figure 1b). After electrophysiological experiments, brain slices were fixed in 4% paraformaldehyde before undergoing CUBIC tissue clearing^51^ and immunohistochemistry to identify pα-syn aggregates within the SNc. As shown in Figure 1c, slices were also stained for tyrosine hydroxylase (TH; green), to identify dopaminergic neurons, and Aldh1a1 (red), a marker of ventral-tier SNc neurons^19,22^. We found a time-dependent increase in pα-syn inclusions in PFF-injected mice (Figure 1d). At 6 weeks post-injection, there was a small, but significant increase in the density of pα-syn inclusions in the SNc in PFF- versus monomer-injected mice. By 12 weeks post-injection, we found a large and significant increase in pα-Syn inclusion number in slices from PFF- versus monomer-injected mice. We also observe that the somas containing pα-syn aggregates lose both TH and Aldh1a1 expression, consistent with previous reports^42,52^

**Figure 1.**
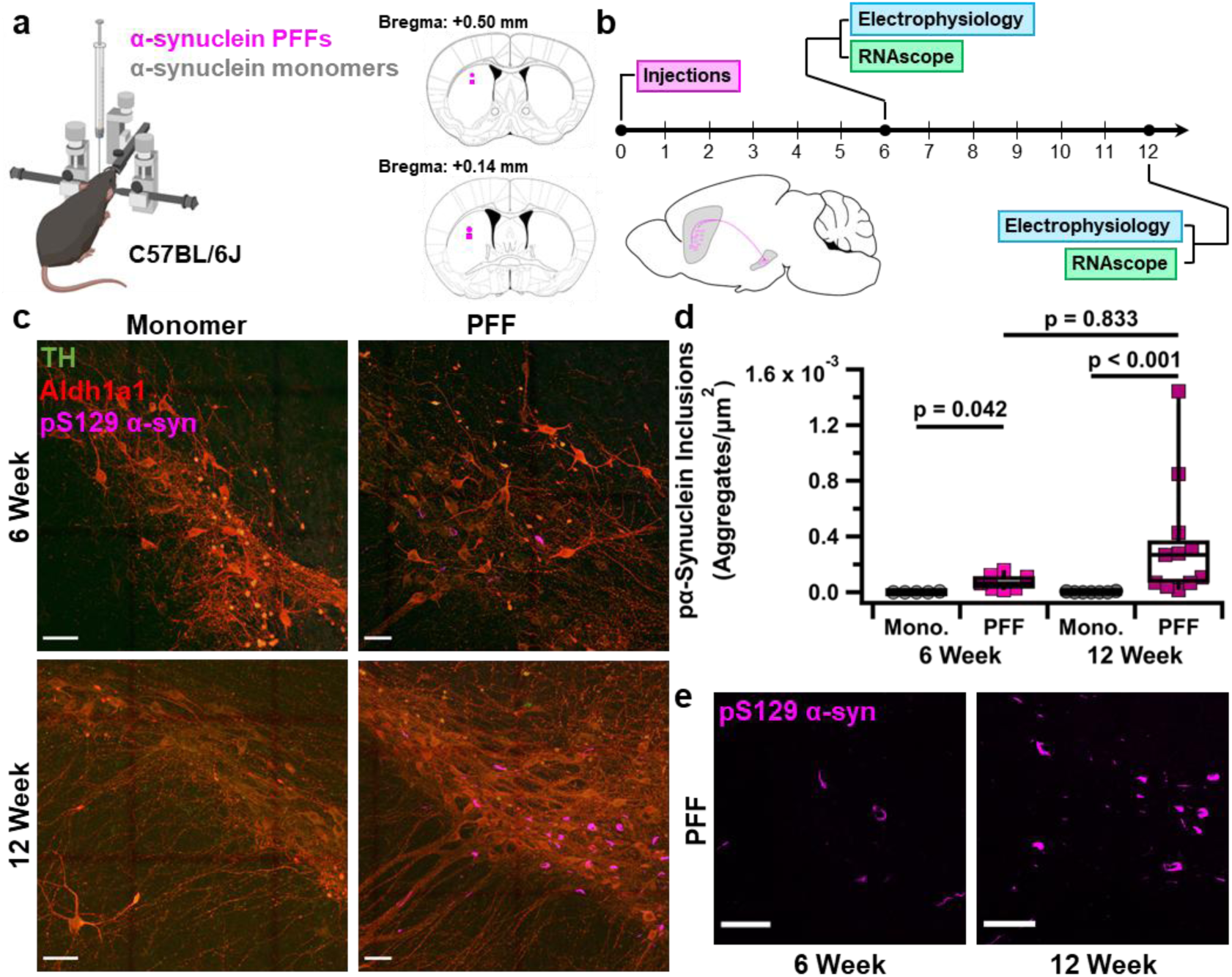
An α-synuclein preformed fibril rodent model of Parkinson’s disease. a, Schematic of experimental design, including atlas images of the dorsolateral striatum indicating α-synuclein injection coordinates. b, Experimental timeline. c, Representative images of the SNc of 6-week monomer (top left), 6-week PFF (top right), 12-week monomer (bottom left), and 12-week PFF (bottom right) injected mice. Scale bars = 50 µm. d, Phosphorylated α-synuclein density is significantly greater in PFF-injected mice vs monomer controls at 6- and 12-weeks post-injection. [6 Week Mono.: 0 (2.83e-06), n = 5 vs PFF: 6.47e-05 (9.74e-05) aggregates/µm2, n = 7; p = 0.042; 12 Week Mono. 0 (5.49e-06), n = 7 vs PFF: 2.68e-04 (3.59e-04) aggregates/µm2, n = 11; p < 0.001; Kruskal-Wallis test followed by Dunn’s test] e, Representative images of phospho-α-synuclein inclusions in the SNc of PFF-injected mice at 6 (left) and 12 (right) weeks post-injection (close up of images in panel c).

### SNc neurons of animals injected with PFFs show decreased dendritic complexity

While SNc axonal degeneration has been widely documented in multiple PD models^54–57^, only a handful of studies address somatodendritic morphological alterations^58–62^. These studies generally observe reduced dendritic arborization in more advanced stages of disease progression. For example, SNc dendritic truncation in MitoPark mice emerges at 16-20 weeks of age^63^, at 6 weeks in the MCI-park mouse^59^, and is apparent in 12-13 month old rats over-expressing α-synuclein, but not at 5 months old^64^. However, SNc dendritic arborization has not yet been assessed using the α-synuclein PFF mouse model. To analyze the effects of α-synuclein PFFs on SNc neuron morphology, we performed electrophysiology experiments using an internal recording solution containing Neurobiotin. After recording, in addition to antibodies targeting TH, Aldh1a1, and pα-syn, slices were stained using a DyLight 405-conjugated streptavidin to visualize Neurobiotin-filled neurons and their processes. Using Neurolucida software, we reconstructed the somas and dendrites of filled cells and analyzed dendritic morphology (Figure 2a). We performed a Sholl analysis to measure dendritic complexity, as the number of intersections of dendrites with concentric circles in increments of 10 μm from the soma. In comparing cells from monomer- and PFF-injected animals of all cell types, we found a trend toward decrease in dendritic complexity at 6-weeks post-injection in PFF-injected animals as measured by the area under the curve (AUC) of the Sholl analysis (Figure 2b & c). However, at 12 weeks post-injection, there was no difference in dendritic complexity (Figure 2d & e). Therefore, PFF injection into the striatum may transiently reduce SNc dendritic branching before extensive midbrain α-synuclein aggregation has occurred.

**Figure 2.**
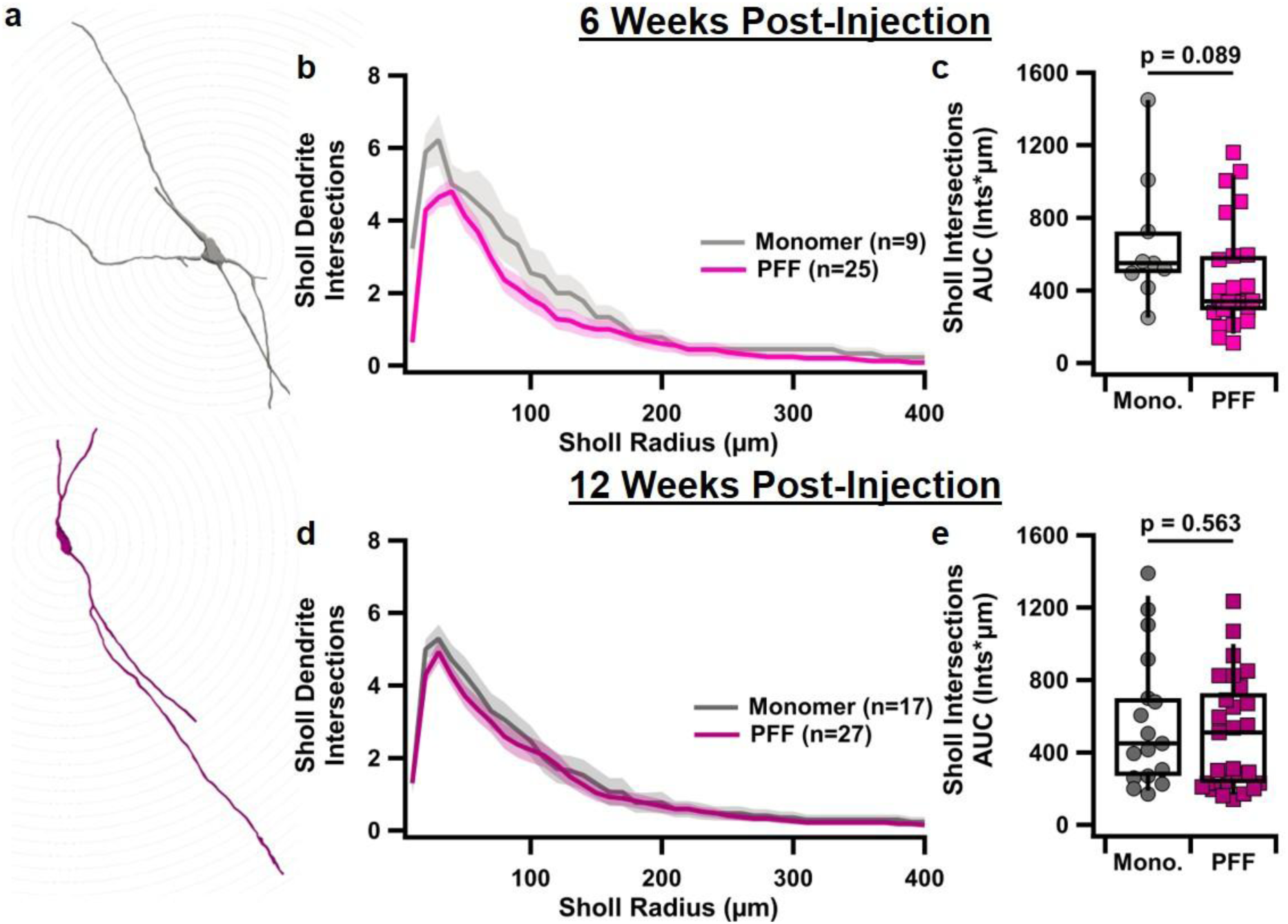
SNc neurons show decreased dendritic complexity at 6 weeks post-PFF injection. a, Representative image of reconstructed SNc neuron from 12-week monomer- (top) and 12-week PFF-(bottom) injected mice. b, Graph showing Sholl intersections as a function of the distance from the soma (Sholl radius) in monomer and PFF groups at 6 weeks post-injection. c, Box plot comparing the area under the curve of Sholl intersections. [Mono.: 550 (413) Intersections*µm, n = 9 vs PFF: 340 (308) Intersections*µm, n = 25; p = 0.026, Wilcoxon rank-sum test] d & e, Same as b & c, but for 12-weeks post-injection. [Mono.: 450 (543) Intersections*µm, n = 17 vs PFF: 510 (535) Intersections*µm, n = 27; p = 0.563, Wilcoxon rank-sum test]

### α-synuclein PFF injection selectively narrows action potential width in vulnerable SNc neurons

Prior to significant neural degeneration, parkinsonian pathology can disrupt the function of SNc dopaminergic neurons by altering their activity levels and firing patterns. Previous studies have shown decreases in spontaneous activity in mitochondria-mediated PD models^59,65^, but increases in activity in α-synuclein overexpression models^16,64^. These studies all consider the SNc as a homogenous structure. However, the SNc contains both vulnerable and resilient populations that may be differentially affected by α-synuclein. Using the electrophysiological signature of vulnerable SNc neurons, a hyperpolarization-induced after-depolarization (ADP)^28^, we separately analyzed spontaneous firing patterns and action potential characteristics of vulnerable (ADP-expressing) and resilient (ADP-lacking) dopaminergic SNc neurons from monomer- and PFF-injected mice. Using *ex vivo* whole-cell electrophysiology, we found that most of the spontaneous action potential characteristics were unchanged by PFF injection. However, in vulnerable SNc neurons, there was a significant decrease in the average half-width of action potentials from PFF-injected mice at 12 weeks as compared to their monomer-injected counterparts (Figure 3c). These cells did not show significant differences in any other measured parameter, including firing rate and action potential height, at 12 weeks. There were no differences in action potential frequency or other characteristics at the 6-week time point (Figure 3a-d). Resilient cells showed no significant differences between monomer-and PFF-injected conditions in firing rate or action potential half-width or height at 6 weeks or 12 weeks post-injection (Figure 3e-f). Thus, narrower action potentials progressively develop only in vulnerable SNc neurons. Previous work in AAV or PFF models of α-synuclein seeding at 4 weeks post-injection found increases in firing frequency in the AAV model, but no changes in action potential half-width^16^, while work in the Mito-Park mouse found progressive narrowing of action potential width with no change in firing frequency in SNc dopaminergic neurons^66^, suggesting a potential common underlying channel disruption mechanism in models with longer term α-synuclein exposure.

**Figure 3.**
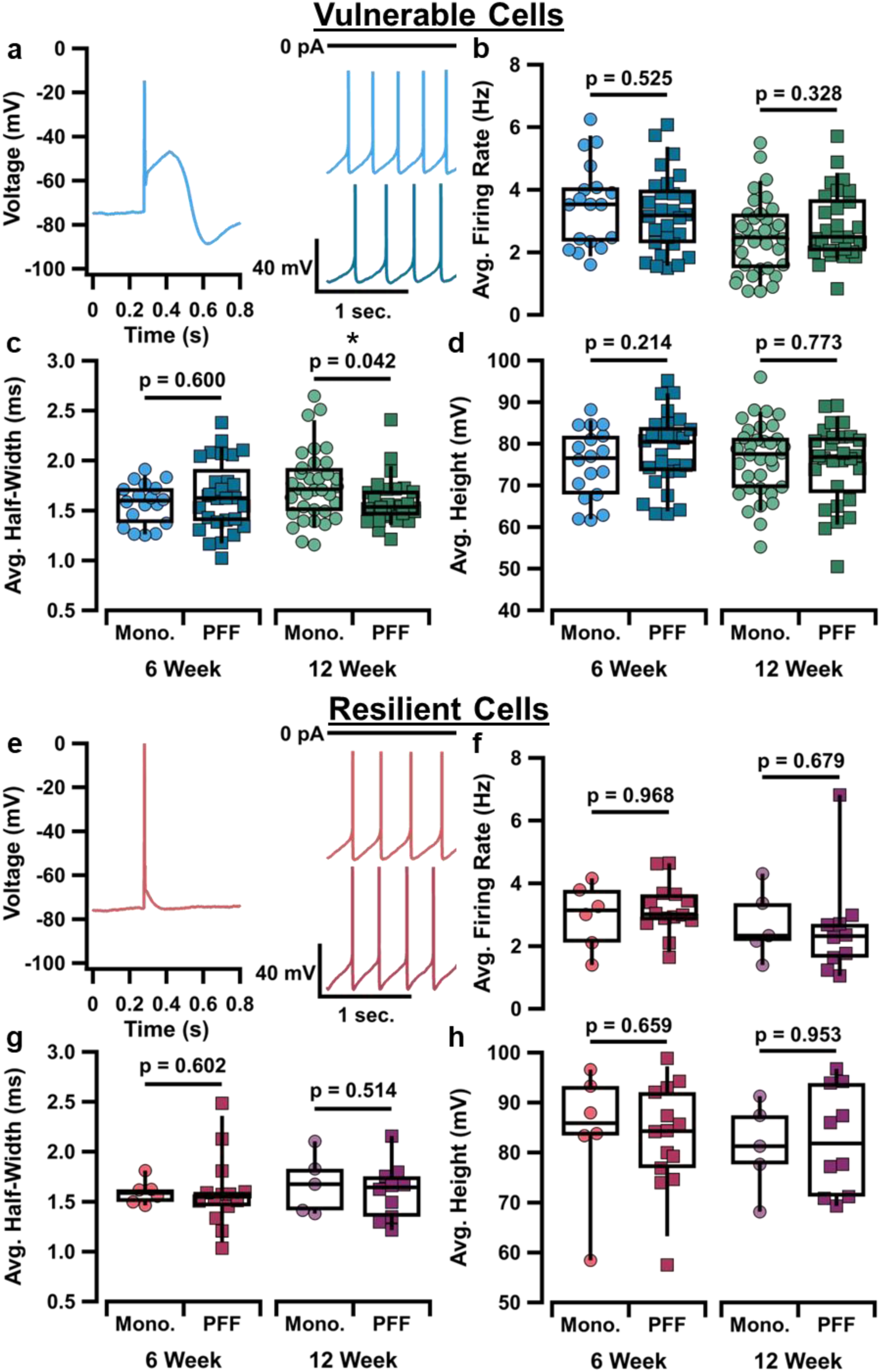
Selective narrowing of AP width in vulnerable SNc dopaminergic neurons in the α-syn PFF model. a, Sample traces of the ADP (left) and tonic firing patterns in vulnerable cells of monomer (top right) and PFF (bottom right) injected mice. b, Box plots comparing 6-week (left) and 12-week (right) measures of average firing rate between monomer and PFF groups in vulnerable cells of the SNc. [6 Week Mono.: 3.53 (1.96) Hz, n = 18 vs PFF: 3.18 (1.78) Hz, n = 28, p = 0.525; 12 Week Mono.: 2.47 (1.76) Hz, n = 34 vs PFF: 2.49 (1.72) Hz, n = 28, p = 0.328; Wilcoxon rank sum tests] c, Box plots comparing 6-week (left) and 12-week (right) measures of average action potential half-width between monomer and PFF groups in vulnerable cells of the SNc. [6 Week Mono.: 1.60 (0.37) ms, n = 18 vs PFF: 1.62 (0.59) ms, n = 28, p = 0.600; 12 Week Mono.: 1.71 (0.45) ms, n = 34 vs PFF: 1.54 (0.27) ms, n = 28; p = 0.042; Wilcoxon rank sum tests] d, Box plots comparing 6-week (left) and 12-week (right) measures of average action potential height between monomer and PFF groups in vulnerable cells of the SNc. [6 Week Mono.: 76.5 (15.0) mV, n = 18 vs PFF: 80.5 (11.0) mV, n = 28, p = 0.214; 12 Week Mono.: 77.5 (12.6) mV, n = 34 vs PFF: 76.8 (14.3) mV, n = 28, p = 0.773; Wilcoxon rank sum tests] e, Sample traces of the absent ADP (left) and tonic firing patterns in vulnerable cells of monomer (top right) and PFF (bottom right) injected mice. f, Box plots comparing 6-week (left) and 12-week (right) measures of average firing rate between monomer and PFF groups in resilient cells of the SNc. [6 Week Mono.: 3.14 (1.95) Hz, n = 6 vs PFF: 3.01 (0.86) Hz, n = 14, p = 0.968; 12 Week Mono.: 2.33 (2.06) Hz, n = 5 vs PFF: 2.32 (1.25) Hz, n = 10, p = 0.679; Wilcoxon rank sum tests] g, Box plots comparing 6-week (left) and 12-week (right) measures of average action potential half-width between monomer and PFF groups in resilient cells of the SNc. [6 Week Mono.: 1.59 (0.18) ms, n = 6 vs PFF: 1.55 (0.23) ms, n = 14, p = 0.602; 12 Week Mono.: 1.68 (0.57) ms, n = 5 vs PFF: 1.64 (0.42) ms, n = 10, p = 0.514; Wilcoxon rank sum tests] h, Box plots comparing 6-week (left) and 12-week (right) measures of average action potential height between monomer and PFF groups in resilient cells of the SNc. [6 Week Mono.: 85.9 (17.0) mV, n = 6 vs PFF: 84.3 (16.0) mV, n = 14, p = 0.659; 12 Week Mono.: 81.3 (16.5) mV, n = 5 vs PFF: 81.8 (22.9) mV, n = 10, p = 0.953; Wilcoxon rank sum tests]

### α-synuclein PFF injection differentially alters vulnerable and resilient SNc neuron excitability

Previous work in Mito-Park and MCI-Park mice show that mitochondrial damage decreases SNc neuron capacitance and increases input resistance^59,66^. Input resistance, which relates to cellular excitability, is increased in α-synuclein overexpressing rats^64^. In mice injected with an α-synuclein*-*expressing AAV or PFFs, no change in capacitance was observed 4 weeks post injection. Further, previous studies did not compare electrophysiological excitability characteristics across distinct subpopulations within the SNc. To examine the effects of α-synuclein PFFs on excitability in vulnerable and resilient SNc cells, we ran a current clamp input-frequency curve holding the cell at approximately -65 mV to inhibit firing prior to depolarizing current injection. Figure 4a shows increasing amounts of depolarizing current inducing increasing firing rates, until reaching depolarization block, at which point voltage-gated sodium channels are unable to return to their resting state and action potentials cannot be induced^67^. In vulnerable SNc neurons, there was no significant difference in the average firing frequency at +10 pA or +100 pA current injection between monomer and PFF groups at 6- or 12-weeks post-injection (Figure 4c & d). Additionally, vulnerable cells from PFF-injected animals do not reach depolarization block any earlier than monomers, as measured at +100 pA injected current (Figure 4b). In resilient SNc neurons, we found a trend toward increased action potential frequency with +100 pA current injection at both the 6-week and 12-week time points. Interestingly, none of the resilient cells from the PFF-injected groups reached depolarization block by the +100-pA current step at 6- or 12- weeks post-injection, a significant difference from the monomer group at 12, but not 6, weeks (Figure 4g). Interestingly, we found no differences in input resistance or membrane capacitance in either vulnerable or resilient SNc neurons at either time point (Supplemental Figure 1). Similarly, we measured the current-voltage relationship of both SNc cell types using a current clamp protocol that injects hyperpolarizing (-150 pA) to depolarizing (+200 pA) currents in a step-wise manner. We found no difference in the change in membrane potential at any current injection step between monomer- and PFF-injected groups at either timepoint in vulnerable or resilient cells (Supplemental Figure 2). Therefore, PFF injection differentially alters the excitability of vulnerable vs resilient SNc neurons, increasing the excitability and the ability to sustain action potential firing selectively in the resilient neurons.

**Figure 4.**
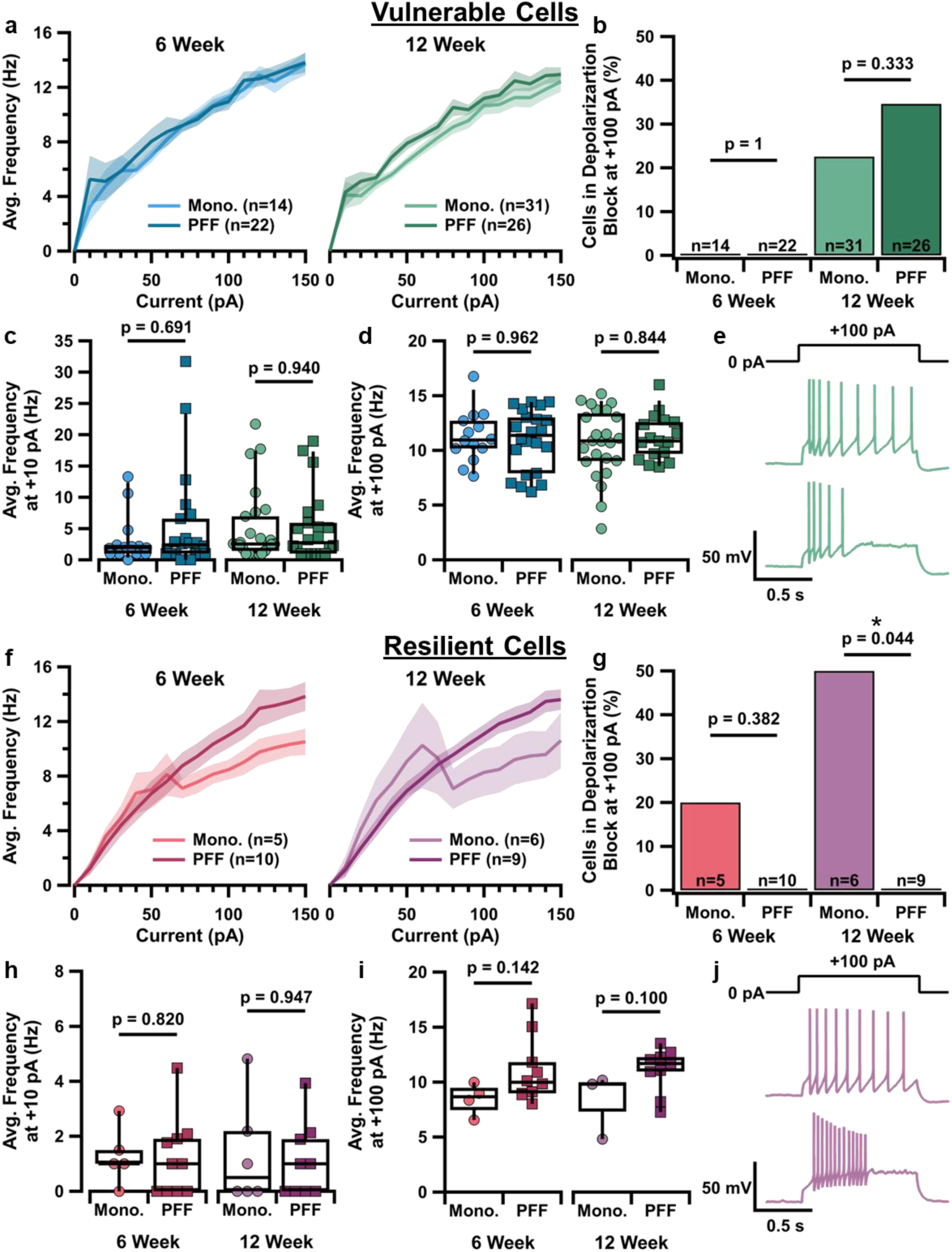
Excitability of SNc subpopulations in an α-syn PFF model. a, Graph of current-frequency relationship in vulnerable cells of monomer and PFF groups at 6 weeks (left) and 12 weeks (right) post-injection. b, Bar graph of percentage of vulnerable cells in depolarization block at +100 pA current injection step in monomer and PFF groups at 6 weeks (left) and 12 weeks (right) post-injection. [6 Week Mono.: 0/14 cells (0%) vs PFF: 0/22 cells (0%), p = 1; 12 Week Mono.: 7/31 cells (22.58%) vs PFF: 9/26 cells (34.62%), p = 0.333; Fisher’s exact test] c, Box plots comparing average firing rate at +10 pA current injection step in vulnerable cells between monomer and PFF groups at 6-weeks (left) and 12-weeks (right) post-injection. [6 Week Mono.: 2.06 (1.61) Hz, n = 16 vs PFF: 2.71 (5.93) Hz, n = 21, p = 0.691; 12 Week Mono.: 2.53 (6.07) Hz, n = 25 vs PFF: 2.74 (5.35) Hz, n = 22, p = 0.940; Wilcoxon rank sum tests] d, Box plots comparing average firing rate at +100 pA current injection step in vulnerable cells between monomer and PFF groups at 6-weeks (left) and 12-weeks (right) post-injection. [6 Week Mono.: 11.00 (2.79) Hz, n = 18 vs PFF: 11.40 (4.56) Hz, n = 24, p = 0.962; 12 Week Mono.: 10.90 (4.49) Hz, n = 24 vs PFF: 10.90 (3.30) Hz, n = 17, p = 0.844; Wilcoxon rank sum tests] e, Sample traces of current-clamp protocol at +100 pA injection step from a resilient cell firing regularly (top) and a cell in depolarization block (bottom) from a 12-week monomer-injected mouse. f, Graph of current-frequency relationship in resilient cells of monomer and PFF groups at 6 weeks (left) and 12 weeks (right) post-injection. g, Bar graph of percentage of resilient cells in depolarization block at +100 pA current injection step in monomer and PFF groups at 6 weeks (left) and 12 weeks (right) post-injection. [6 Week Mono.: 1/5 cells (20%) vs PFF: 0/10 cells (0%), p = 0.382; 12 Week Mono.: 3/6 cells (50%) vs PFF: 0/9 cells (0%), p = 0.044; Fisher’s exact test] h, Box plots comparing average firing rate at +10 pA current injection step in resilient cells between monomer and PFF groups at 6-weeks (left) and 12-weeks (right) post-injection. [6 Week Mono.: 1.00 (1.85) Hz, n = 6 vs PFF: 1.00 (1.91) Hz, n = 11, p = 0.820; 12 Week Mono.: 0.50 (2.85) Hz, n = 6 vs PFF: 1.00 (2.01) Hz, n = 9, p = 0.947; Wilcoxon rank sum tests] i, Box plots comparing average firing rate at +100 pA current injection step in resilient cells between monomer and PFF groups at 6-weeks (left) and 12-weeks (right) post-injection. [6 Week Mono.: 8.35 (2.54) Hz, n = 5 vs PFF: 9.85 (2.82) Hz, n = 11, p = 0.142; 12 Week Mono.: 9.81 (5.38) Hz, n = 3 vs PFF: 11.70 (2.91) Hz, n = 9, p = 0.100; Wilcoxon rank sum tests] j, Sample traces of current-clamp protocol at +100 pA injection step from a resilient cell firing regularly (top) and a cell in depolarization block (bottom) from a 12-week monomer-injected mouse.

### α-synuclein PFF injection does not alter excitatory synaptic input onto SNc subpopulations

α-synuclein contributes to synaptic vesicle release^68^. Misfolded α-synuclein disrupts synaptic plasticity in the striatum^69^ and reduces spine density in the hippocampus^70^ and cortex^46^, and functional α-synuclein is necessary for somatodendritic dopamine release^71^. However, it is not known whether misfolded α-synuclein alters the excitatory synaptic input onto the vulnerable and resilient SNc dopaminergic subpopulations. We recorded spontaneous excitatory post-synaptic currents (sEPSCs) in SNc neurons using a voltage-clamp protocol holding the membrane potential at -70 mV. Events were detected by the Neuromatic plugin for Igor and visually confirmed by the reviewer as previously^72^. We measured these excitatory inputs by event amplitude and frequency, averaging all events per cell. Cells that did not have any events were excluded from the compiled amplitude data, but were included in the frequency measurements as 0 Hz. Interestingly, we found a significant difference in sEPSC frequency between vulnerable and resilient cells across control (monomer) groups, with resilient neurons having significantly higher sEPSC frequencies (Supplemental Figure 3). However, PFF injection did not change the sEPSC amplitude or frequency in either vulnerable or resilient neurons at either time point (Figure 5). These results show that intra-striatal PFFs do not alter excitatory synaptic inputs to SNc neurons 6 or 12 weeks after injection.

**Figure 5.**
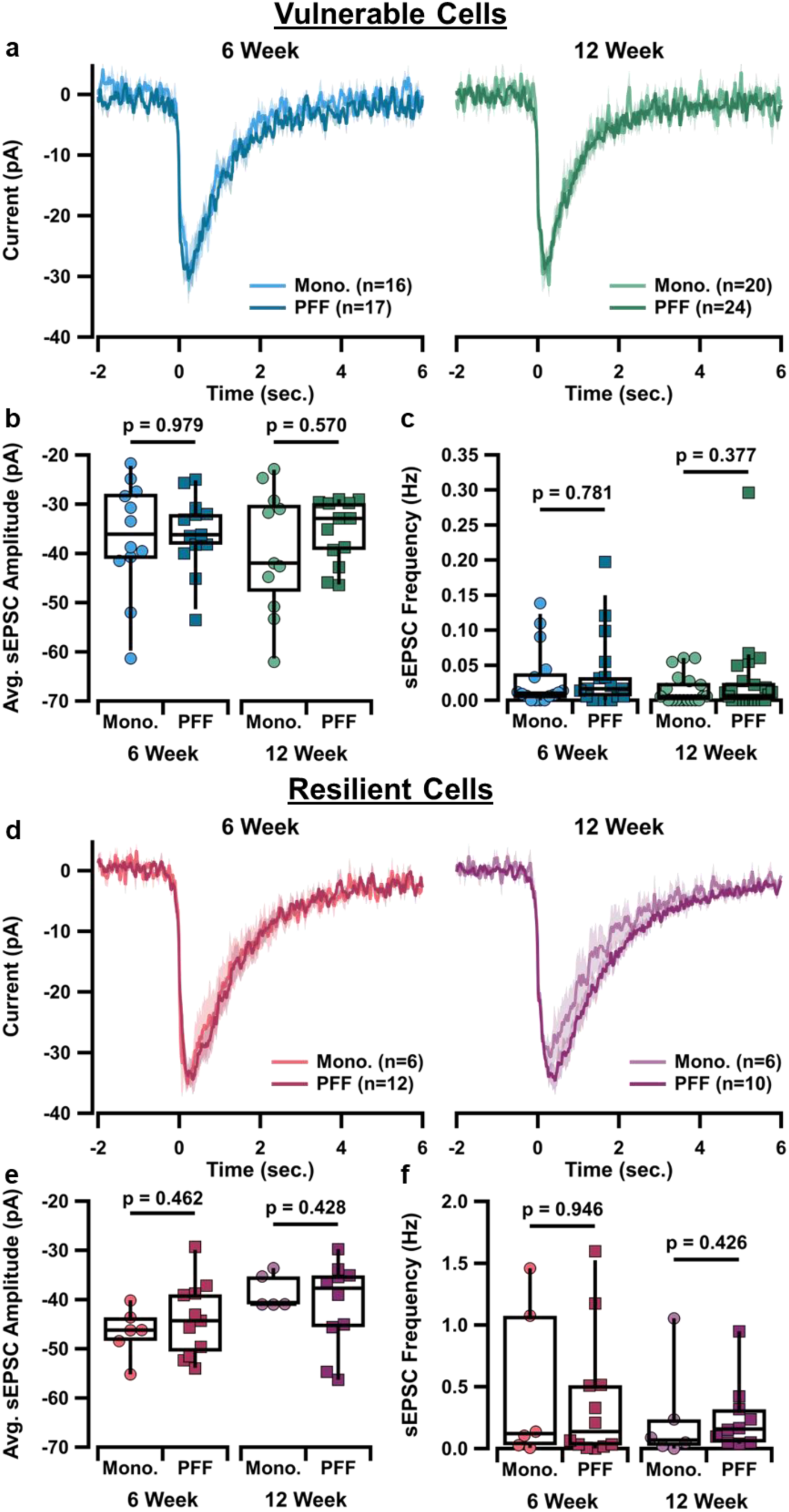
Spontaneous excitatory post-synaptic inputs to SNc neurons in an α-syn PFF model. a, Graph of sEPSC averages from vulnerable cells of monomer and PFF groups at 6 weeks (left) and 12 weeks (right) post-injection. b, Box plots comparing average sEPSC amplitude in vulnerable cells between monomer and PFF groups at 6-weeks (left) and 12-weeks (right) post-injection. [6 Week Mono.: -36.1(13.6) pA, n = 12 vs PFF: -36.2 (7.7) pA, n = 13, p = 0.979; 12 Week Mono.: -42.0 (21.5) pA, n = 11 vs PFF: -32.9 (11.4) pA, n = 13, p = 0.570; Wilcoxon rank sum tests] c, Box plots comparing sEPSC frequency in vulnerable cells between monomer and PFF groups at 6-weeks (left) and 12-weeks (right) post-injection. [6 Week Mono.: 0.010 (0.040) Hz, n = 16 vs PFF: 0.016 (0.041) Hz, n = 17, p = 0.781; 12 Week Mono.: 0.003 (0.023) Hz, n = 18 vs PFF: 0.011 (0.038) Hz, n = 21, p = 0.377; Wilcoxon rank sum tests] d, Graph of sEPSC averages from resilient cells of monomer and PFF groups at 6 weeks (left) and 12 weeks (right) post-injection. e, Box plots comparing average sEPSC amplitude in resilient cells between monomer and PFF groups at 6-weeks (left) and 12-weeks (right) post-injection. [6 Week Mono.: -46.2 (7.3) pA, n = 6 vs PFF: -44.3 (12.7) pA, n = 11, p = 0.462; 12 Week Mono.: -41.0 (6.5) pA, n = 5 vs PFF: -37.7 (13.1) pA, n = 10, p = 0.428; Wilcoxon rank sum tests] f, Box plots comparing sEPSC frequency in resilient cells between monomer and PFF groups at 6-weeks (left) and 12-weeks (right) post-injection. [6 Week Mono.: 0.121 (1.148) Hz, n = 6 vs PFF: 0.137 (0.493) Hz, n = 12, p = 0.946; 12 Week Mono.: 0.069 (0.425) Hz, n = 6 vs PFF: 0.158 (0.295) Hz, n = 10, p = 0.426; Wilcoxon rank sum tests]

### α-synuclein aggregation reduces T-type calcium currents in vulnerable SNc neurons

The vulnerable, ventral-tier neurons of the SNc have larger T-type calcium currents that mediate their rebound properties^73^ and characteristic ADP^28^. Using a voltage-clamp ramp protocol, we measured T-type calcium currents by gradually increasing the holding potential from -90 mV to -50 mV over 1 second. Because the T-type currents often had multiple peaks, we calculated the area under the curve (AUC) between -75 mV and -55 mV as the measure of TTCC current density (Figure 6a & e). Leak current was subtracted with a linear fit. In vulnerable SNc cells we found that T-type calcium current is significantly decreased in PFF-injected animals as compared to monomer-injected controls at the 12-week timepoint, but not at the 6-week timepoint (Figure 6b). Because vulnerable SNc neurons show more T-type mediated rebound activity^28,74^, we evaluated the effect of PFF injection on SNc rebound characteristics in vulnerable neurons. Surprisingly, we found no significant difference in rebound slope between monomer and PFF conditions at 6- or 12-week timepoints (Figure 6c). Further, there was no significant difference in size of the ADP at either timepoint.

**Figure 6.**
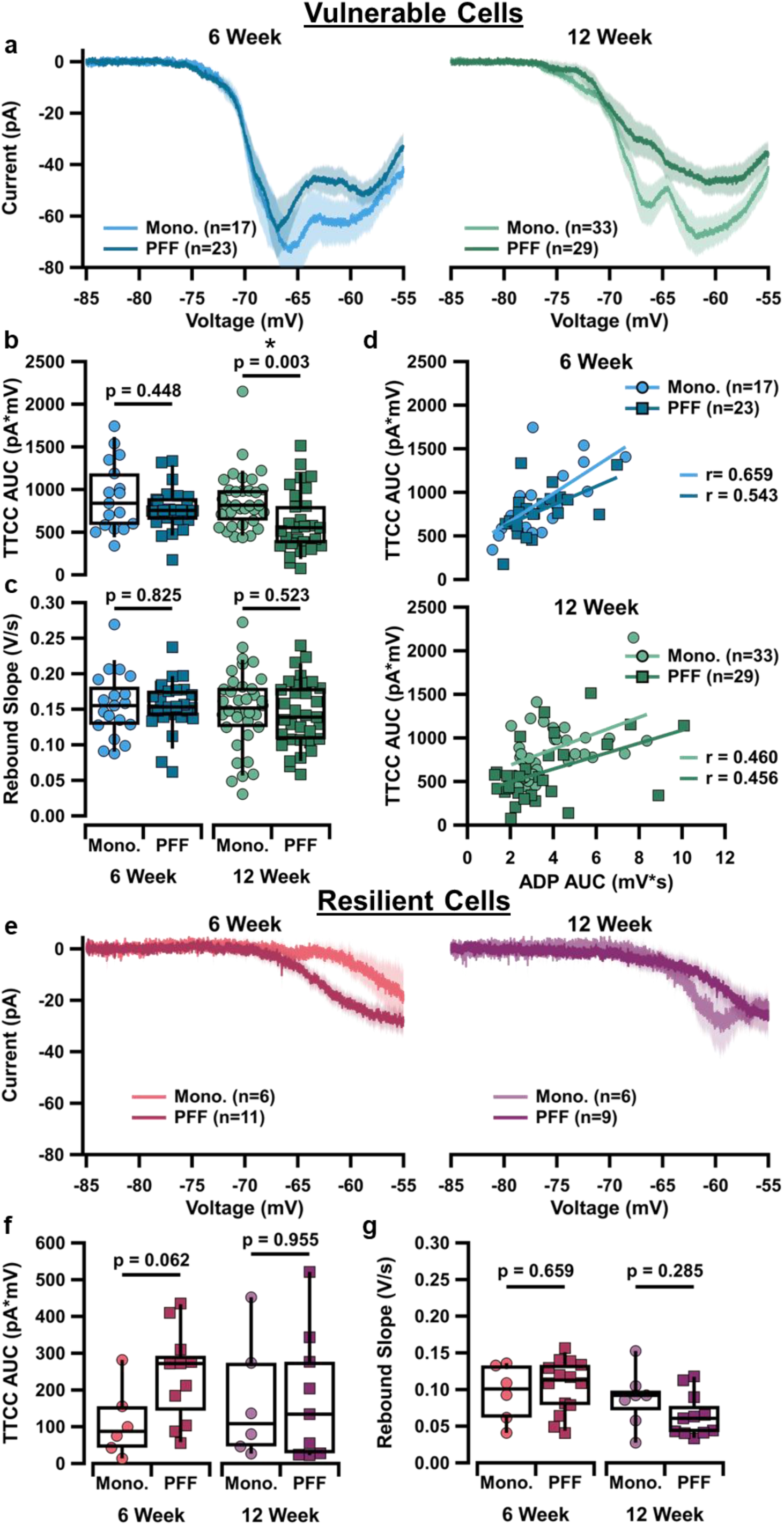
T-type calcium current activity in SNc neurons in an α-syn PFF model. a, Graph of average T-type calcium current in vulnerable cells of monomer and PFF groups at 6 weeks (left) and 12 weeks (right) post-injection. b, Box plots comparing area under the curve (AUC) of T-type calcium current in vulnerable cells between monomer and PFF groups at 6-weeks (left) and 12-weeks (right) post-injection. [6 Week Mono.: 839 (707) pA*mV, n = 17 vs PFF: 755 (276) pA*mV, n = 23, p = 0.448; 12 Week Mono.: 815 (388) pA*mV, n = 33 vs PFF: 552 (512) pA*mV, n = 29, p = 0.003; Wilcoxon rank sum tests] c, Box plots comparing rebound slope in vulnerable cells between monomer and PFF groups at 6-weeks (left) and 12-weeks (right) post-injection. [6 Week Mono.: 0.155 (0.064) V/s, n = 19 vs PFF: 0.153 (0.038) V/s, n = 27, p = 0.825; 12 Week Mono.: 0.152 (0.056) V/s, n = 36 vs PFF: 0.139 (0.075) V/s, n = 33, p = 0.523; Wilcoxon rank sum tests] d, Graph of T-type calcium AUC relative to AUC of ADP from vulnerable cells of monomer and PFF groups at 6 weeks (top) and 12 weeks (bottom) post-injection. e, Graph of average T-type calcium current in resilient cells of monomer and PFF groups at 6 weeks (left) and 12 weeks (right) post-injection. f, Box plots comparing area under the curve (AUC) of T-type calcium current in resilient cells between monomer and PFF groups at 6-weeks (left) and 12-weeks (right) post-injection. [6 Week Mono.: 87.5 (151.3) pA*mV, n = 6 vs PFF: 272 (204) pA*mV, n = 12, p = 0.062; 12 Week Mono.: 108.0 (277.4) pA*mV, n = 6 vs PFF: 134 (284.2) pA*mV, n = 9; p = 0.955; Wilcoxon rank sum tests] g, Box plots comparing rebound slope in resilient cells between monomer and PFF groups at 6-weeks (left) and 12-weeks (right) post-injection. [6 Week Mono.: 0.101 (0.078) V/s, n = 6 vs PFF: 0.113 (0.060) V/s, n = 14, p = 0.659; 12 Week Mono.: 0.092 (0.047) V/s, n = 7 vs PFF: 0.061 (0.049) V/s, n = 11, p = 0.285; Wilcoxon rank sum tests]

The rebound slope and ADP characteristics are both a function of TTCC activity. We found a direct correlation between TTCC AUC measured in voltage-clamp and the AUC of the ADP at 6- and 12-weeks post-injection in monomer- and PFF-injected conditions (Figure 8d). This correlation was the strongest in the 6-week monomer group. Though rebound slope and the ADP require TTCC activity, there are other factors that confound these as a direct measure of TTCC current density. These data indicate that α-synuclein aggregation in the SNc inhibits TTCC, but suggest that additional mechanisms contribute to rebound slope.

In resilient, non-ADP SNc neurons, our recordings show much less TTCC activity than in vulnerable, ADP-expressing SNc neurons under control (monomer) conditions, consistent with previous findings^28^. Unlike the vulnerable SNc neurons, we saw no effect of PFFs on TTCC activity in resilient SNc neurons. There is no change in TTCC density, as measured by the AUC, in PFF-injected animals at 6- or 12-weeks post-injection (Figure 6f). As in vulnerable SNc neurons, there is no difference in rebound slope between monomer- and PFF-injected resilient SNc neurons at 6- or 12-weeks post-injection (Figure 6g). These data suggest that the resilient cells are minimally affected by any potential TTCC disruptions resulting from PFF injection.

### Hyperpolarization-activated cation channel activity is not altered by the presence of α-synuclein aggregates

Dopamine neurons of the SNc are also characterized by hyperpolarization-activated cation (HCN) channels, which work with TTCCs to enhance rebound activity^28,75–77^. Rats over-expressing α-synuclein show a reduction^78^ or no change^64^ in HCN-mediated current (I_h_) in dopaminergic neurons, while I_h_ was also reduced in the SNc of MCI-Park mice^59^. Because of the observed changes in T-type current without any changes in the measures of rebound, we hypothesized that an increase in I_h_ could counteract the reduction in TTCC current to maintain rebound activity at baseline levels. Using voltage clamp recordings of I_h_ tail currents as previously^74^, we determined the voltage of half-maximum activation (V50). In vulnerable cells, we found no difference in the V50 of I_h_ between monomer and PFF conditions at 6- or 12-week timepoints (Figure 7b). Similarly, in resilient cells, we also found no difference in the V50 of I_h_ at 6 weeks or 12 weeks post-injection between monomer and PFF groups (Figure 7d). Together, these results suggest that changes in I_h_ are not likely compensating for TTCC reduction in measures of rebound.

**Figure 7.**
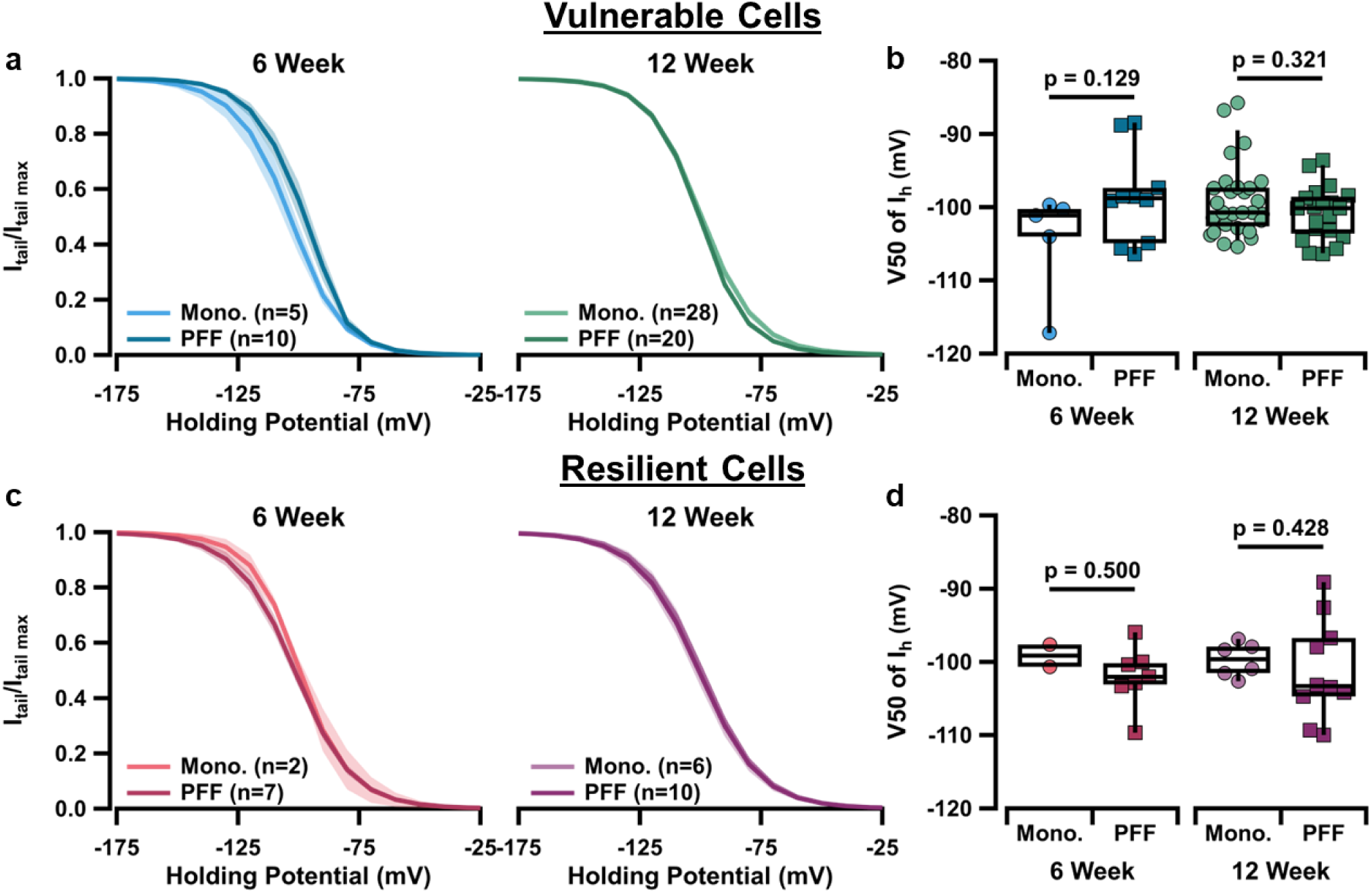
Ih in SNc neurons in an α-syn PFF model. a, Normalized activation curves of Ih tail current in vulnerable cells of monomer and PFF groups at 6 weeks (left) and 12 weeks (right) post-injection. b, Box plots comparing V50 of Ih in vulnerable cells between monomer and PFF groups at 6-weeks (left) and 12-weeks (right) post-injection. [6 Week Mono.: -101.0 (11.0) mV, n = 5 vs PFF: -98.8 (7.6) mV, n = 10, p = 0.129; 12 Week Mono.: -100.7 (5.3) mV, n = 28 vs PFF: -100.1 (5.0) mV, n = 21, p = 0.321; Wilcoxon rank sum tests] c, Normalized activation curves of Ih tail current in resilient cells of monomer and PFF groups at 6 weeks (left) and 12 weeks (right) post-injection. d, Box plots comparing V50 of Ih in resilient cells between monomer and PFF groups at 6-weeks (left) and 12-weeks (right) post-injection. [6 Week Mono.: -99.1 (3.4) mV, n = 2 vs PFF: -99.6 (3.6) mV, n = 7, p = 0.643; 12 Week Mono.: -102.0 (3.3) mV, n = 6 vs PFF: -103.3 (8.0) mV, n = 10, p = 0.813; Wilcoxon rank sum tests]

### T-type calcium channel mRNA expression is not changed by PFF injection

Based on prior research implicating α-synuclein in TTCC dysregulation and our electrophysiology data, we examined potential changes in calcium channel expression using fluorescence *in situ* hybridization (FISH, RNAscope), targeting mRNA for each of the TTCC channel subtypes: CACNA1G (CaV3.1), CACNA1H (CaV3.2), and CACNA1I (CaV3.3). Consistent with previous work^79,80^, we found that CaV3.1 is the most prominent TTCC expressed in SNc dopaminergic neurons (Figure 8c), while CaV3.2 shows higher expression in the substantia nigra *pars reticulata* (SNr, Figure 8a), with minimal expression of CaV3.3 (Figure 8b). Measuring average raw fluorescence in the SNc for each TTCC mRNA, we found no significant change in any channel type between PFF and monomer injected groups (Figure 8d). These data show that PFFs do not drastically alter TTCC expression in the SNc at the mRNA level, and suggests that the reduction in TTCC current observed in this study is due to modifications of TTCCs at the protein level, such as differential phosphorylation or membrane insertion. However, the sample size was not powered to compare 6- and 12-week post-injection time points, and the two have been combined in this figure, leaving open the possibility that TTCC mRNA could have time-sensitive alterations not detectable in this experiment.

**Figure 8.**
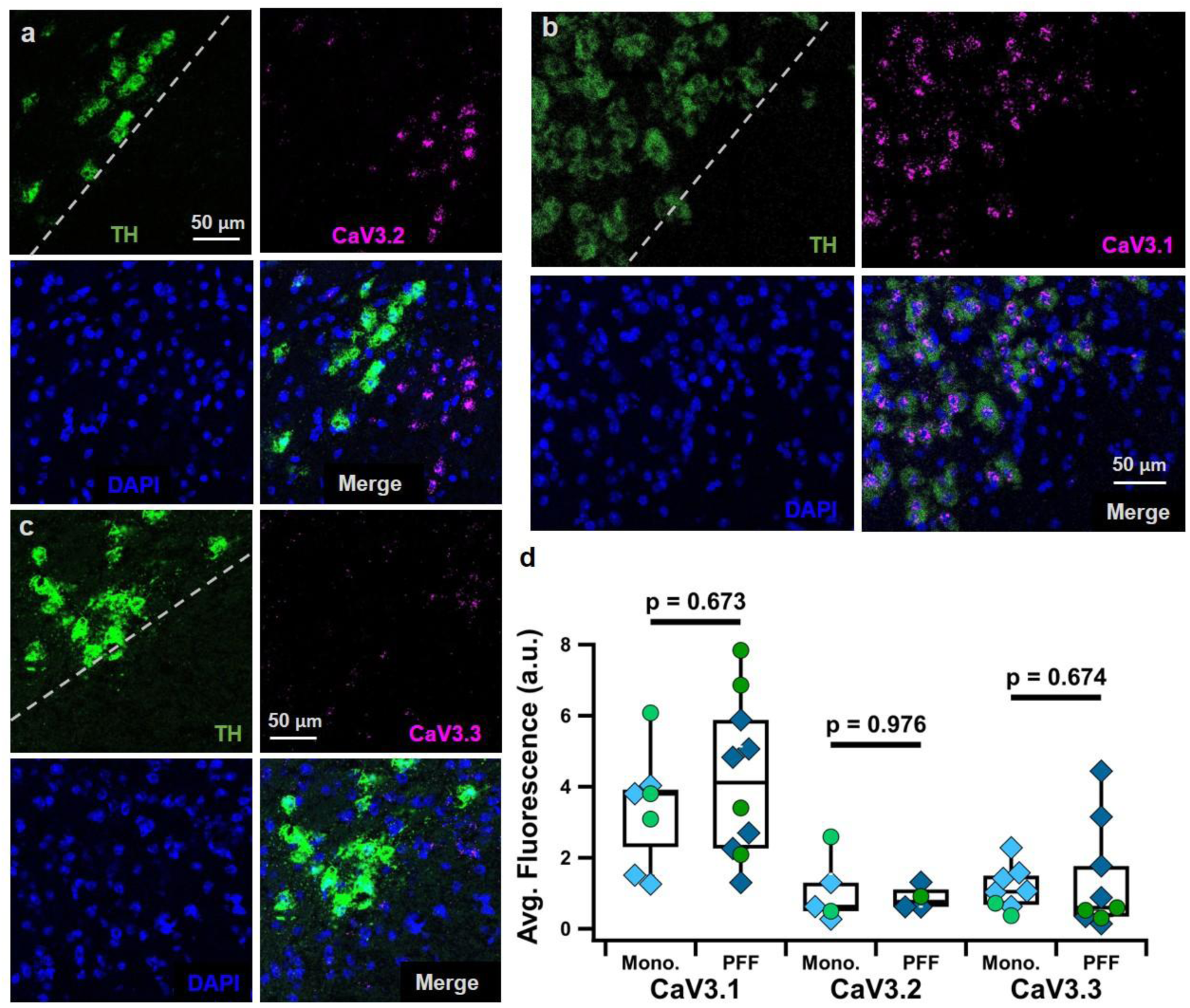
RNAscope analysis of T-type calcium channel mRNA expression in SNc neurons in an α-syn PFF model. a, Sample RNAscope images of CACNA1H (CaV 3.2), dotted line in TH image represents the border between the SNc and substantia nigra pars reticulata (SNr). b. Same as a, but for CACNA1I (CaV 3.3) c, Same as a, but for CACNA1G (CaV 3.1). d, Box plots comparing average fluorescence of CACNA1G (CaV3.1), CACNA1H (CaV3.2), and CACNA1I (CaV3.3) expression in the SNc of monomer- vs PFF-injected animals (CaV3.1 Mono. 3.8 (2.5) a.u., n (slices) = 7 vs PFF 4.1 (3.6) a.u., n = 10, p = 0.673; CaV3.2 Mono. 0.6 (1.6) a.u., n = 5 vs PFF 0.8 (0.5) a.u., n = 4, p = 0.976; CaV3.3 Mono. 1.0 (0.8) a.u., n = 8 vs PFF 0.6 (2.1) a.u., n = 9, p = 0.674). Blue diamonds show 6 week post-injection timepoint, while green circles show 12 week post-injection time point.

## Discussion

Accumulation of Lewy Bodies in dopaminergic neurons of the SNc is a key feature of PD pathology. These aggregates of α-synuclein protein are toxic to neurons and contribute to their eventual dysfunction and degeneration. However, little is known about the effects of α-synuclein on subpopulations of SNc dopaminergic neurons in early stages of disease progression. Here, we found changes in TTCC function of SNc neurons from animals injected with α-synuclein PFFs that provide insight to selective dopaminergic vulnerability in PD and reveal complex interactions between α-synuclein and TTCCs.

We found that α-synuclein burden in the SNc differentially affects the vulnerable and resilient subpopulations. We found a significant decrease in TTCC current from PFF-injected mice in the vulnerable, but not resilient SNc neurons. Importantly, high levels of TTCC current are a defining characteristics of the vulnerable SNc neurons, possibly contributing to their vulnerability by causing extensive dendritic calcium influx in their dendrites^28^. Reduction in TTCC current may actually be a protective response to α-synuclein accumulation, as pharmacological blockade of TTCCs shows promise as a protective treatment in multiple models of PD^39,81,82^. This selective decrease in T-type calcium current could be caused by several factors. 1. α-Synuclein accumulation could downregulate channel expression at the mRNA or protein level. 2. α-Synuclein accumulation could alter the activity of TTCCs through phosphorylation or a change in calcium channel subunits. 3. Reduced TTCC current could be a consequence of dendritic pruning, as TTCCs are prominent on SNc dendrites^28,80^. 4. Although we did not observe obvious loss of SNc neurons after PFF injections, it is possible that α-synuclein makes the neurons with the highest level TTCC expression unhealthy, such that whole-cell recording of these neurons is not feasible.

TTCC signaling is important for vulnerable SNc neurons to recover from inhibition and produce rebound activity^73,83^. While reduction in TTCC signaling may reduce the calcium burden on SNc neurons, loss of TTCC signaling could disrupt healthy information processing in these cells. Therefore, it is encouraging that the reduced TTCC current was not accompanied by a reduction in rebound activity. This finding was surprising because TTCCs are one of the strongest contributors to rebound activity in the vulnerable SNc neurons^28,74^ and a previous study found reduced SNc rebound activity in α-synuclein-based mouse models^16^. We hypothesized that an upregulation in I_h_, another current that is strong in vulnerable SNc neurons and contributes to rebound, could counteract the reduction in TTCC current to maintain rebound activity. However, previous work suggests that α-synuclein would decrease, rather than increase I_h_^78^. Ultimately, we found no differences in I_h_ activation characteristics in vulnerable or resilient SNc neurons from PFF-injected mice. Another possibility is that a downregulation of A-type potassium currents, which work to suppress rebound, could counteract TTCC reduction, maintaining rebound activity. Indeed, previous studies show that mutant α-synuclein^84^ and α-synuclein overexpression^64^ both result in a reduction in A-type potassium current. Future work will be needed to determine whether TTCCs and A-type potassium channels are both downregulated in the PFF model.

Vulnerable SNc neurons showed decreased action potential half-width in the PFF condition. This change could also serve a protective role, as the characteristically wide action potentials in SNc neurons are due to calcium channels opening^85^. Thus, a narrower action potential would reduce dendritic calcium influx. It is interesting that work in a completely different type of PD mouse model, the Mito-Park mouse yielded a similar change in action potential width with no change in firing frequency^66^. This suggests a common pathway or compensatory response is at work in both PD models. Interestingly, we did not observe action potential narrowing or a reduction in TTCC current in the resilient SNc population, demonstrating subpopulation-specific physiological effects of α-synuclein burden.

Importantly, the reduction in TTCC and action potential width were time-dependent, developing as the α-synuclein burden increased from 6 to 12 weeks. This progressive change reflects cellular alterations that could be developing in the prodromal phase of PD, prior to significant dopaminergic cell loss and prior to symptom onset. Understanding these early circuit alterations and how physiological changes develop in response to α-synuclein burden is critical for understanding the way that early intervention therapies would work. Blocking T-type channels, for example, could have a different effect early in disease time course compared to later when TTCC currents were significantly reduced.

In conclusion, unilateral PFF injection in the dorsolateral striatum is sufficient to seed α-synuclein pathology in dopamine neurons of the SNc. The subsequent changes in dopamine function appear as decreased dendritic complexity and subtype-specific changes in T-type calcium channel activity. Importantly, these findings reveal that α-synuclein burden does not affect all SNc neurons equally, preferentially altering the calcium currents and action potential shape of the most vulnerable SNc neuronal subtype.

## Methods

### Animal use

All animal handling and procedures were approved by the Animal Care and Use Committee for Georgetown University. Measures were taken to minimize animal distress or discomfort. Experiments were designed to minimize the number of animals used. Wild-type C57BL/6J mice [JAX# 000664] were bred in house. Adult mice of either sex were injected at 9-14 weeks of age (average age: 72 ± 3 days) and euthanized for electrophysiology or RNAscope experiments 6- or 12-weeks post-injection (average age: 139 ± 5 days).

### α-Synuclein pre-formed fibril preparation

Mouse Recombinant Alpha Synuclein Pre-Formed Fibrils (PFFs) (Type 1) and Mouse Recombinant Alpha Synuclein Monomers (Type 1) were purchased from StressMarq at a concentration of 2 mg/mL. Using a probe sonicator, PFFs were sonicated for two, 1-minute cycles of 1 second on 1 second off at 10% amplitude, with a 2-minute wait in between and aliquots were stored at -80°C. Monomers were mixed with a pipette prior to aliquoting and storage. Aliquots were vortexed after thawing immediately before use.

### Stereotactic injections

Stereotactic injections were performed using a Stoelting 51730UD stereotactic frame and 5 µL Hamilton microsyringe (65460-02). Mice were at least 9 weeks old at the time of surgery. Mice were anesthetized with inhaled 5% isoflurane and maintained on 1-3% isoflurane with oxygen at a flow of 1 L/min for the duration of the procedure. Animals were placed on a heating pad within the stereotactic frame, and the skull was stabilized with the ear bars and nose cone. Bupivacaine (5 mg/kg) and carprofen (5 mg/kg) were administered subcutaneously pre-operatively as a local anesthetic and analgesic, respectively. A small incision was made to expose the skull, where bregma and lambda were visualized to level the skull and serve as reference for injection coordinates. After drilling a small hole at the appropriate coordinates, the syringe was positioned accordingly to infuse at a rate of 0.2 µL/min. Mice were given four 1 µL-injections of either α-synuclein monomers (StressMarq: SPR-323) or PFFs (StressMarq: SPR-324) at a concentration of 2 µg/µL in the left dorsolateral striatum (A-P: +0.14, M-L: +2.00, D-V: - 3.00, -2.65 and A-P: +0.50, M-L: +1.80, D-V: -3.00, -2.65). Designated syringes are used for monomer and PFF injections accordingly to eliminate risk of cross-contamination. Once the first infusion at D-V -3.00mm was finished, the syringe was raised to -2.65mm for the second injection. After the second infusion was complete, the syringe was raised to -2.15 mm and allowed to rest for 10 minutes. This process was repeated for the second set of injections. After all injections were completed, the incision was closed using VetBond tissue adhesive and wound clips. Post-operatively, mice were administered buprenorphine extended-release (1.0 mg/kg) and sterile saline (0.2-0.3 mL) subcutaneously for long-lasting analgesia and hydration. Mice were allowed to recover on a heating pad until resuming normal grooming and feeding behavior, and then returned to a clean cage where they were monitored daily for signs of distress or infection.

### *Ex vivo* slice preparation

Mice were anesthetized with inhaled isoflurane and transcardially perfused with an ice-cold, oxygenated, glycerol-based modified artificial cerebrospinal fluid (aCSF) solution containing the following (in millimolar (mΜ)): 198 glycerol, 2.5 KCl, 1.2 NaH_2_PO_4_, 25 NaHCO_3_, 20 HEPES, 10 glucose, 10 MgCl_2_, and 0.5 CaCl_2_. Mice were then decapitated and brains extracted. Coronal midbrain slices (200 µm) containing the substantia nigra region were prepared using a vibratome (Leica VT 1200S) and incubated for 30 minutes in heated (34°C) oxygenated holding aCSF containing (in mΜ): 92 NaCl, 30 NaHCO_3_, 1.2 NaH_2_PO_4_, 2.5 KCl, 35 glucose, 20 HEPES, 2 MgCl_2_, 2 CaCl_2_, 5 Na-ascorbate, 3 Na-pyruvate, and 2 thiourea, as in Evans et al., 2017. Following, slices incubated in their holding chamber at room temperature for at least 30 minutes. For recording, slices were hemisectioned and continuously superfused at ∼2-3 mL/min with warm (34°C), oxygenated extracellular recording solution containing the following (in mΜ): 125 NaCl, 25 NaHCO_3_, 3.5 KCl, 1.25 NaH_2_PO_4_, 10 glucose, 1 MgCl_2_, and 2 CaCl_2_.

### *Ex vivo* slice electrophysiology

Neurons were visualized with a 40x objective using a Prior OpenStand Olympus microscope equipped with a scientific CMOS camera (Hamamatsu ORCA-spark). All recordings were performed in dopaminergic neurons which were targeted by their anatomic location and identified based on various electrophysiological characteristics, such as the firing frequency (<5 Hz) and presence of HCN-mediated sag. Ventral-tier (vulnerable) SNc neurons were identified by the presence of the distinctive ADP^28^.

Whole-cell recordings were made using borosilicate pipettes (2-5 MΩ) pulled with a flaming/brown micropipette puller (Sutter Instruments) and filled with internal recording solution containing (in mΜ): 121.5 KMeSO_3_, 9 NaCl, 9 HEPES, 1.8 MgCl_2_, 14 phosphocreatine, 4 Mg-ATP, 0.3 Na-GTP, 0.1 CaCl_2_, and 0.5 EGTA adjusted to a pH value of ∼7.35 with KOH. The internal solution also contained Neurobiotin (0.1-0.3%) for post-hoc staining of patched cells.

Signals were digitized with an Axon Digidata 1550B interface, amplified by a Multiclamp 700B amplifier, and acquired using Clampex11.2 software (Molecular Devices). Data were sampled in current clamp at 10 kHz and in voltage clamp at 100kHz with filtering at 5 kHz. Data were analyzed using custom procedures in Igor Pro (WaveMetrics) and GraphPad Prism.

### Tissue clearing and immunohistochemistry

After electrophysiology experiments, slices were fixed overnight in 4% paraformaldehyde in phosphate buffer (PB) solution, pH 7.6 at 4°C and then stored in PB solution at 4°C until immunostaining. The CUBIC tissue clearing protocol^86^ was combined with immunohistochemistry, as previously^73,87^. All steps are performed with slices in a 24-well plate, at room temperature, on a rotating platform. Slices were placed in CUBIC Reagent 1 overnight, then washed in PB (3x 1 hour) and placed in blocking solution (0.5% fish gelatin in PB) for 3 hours, before being placed in primary antibodies for 2-3 days. Then, slices were washed again in PB (3x 2 hours) and placed in secondary antibodies for 2-3 days. Lastly, slices were washed in PB (3x 2 hours) and placed in CUBIC Reagent 2 overnight before mounting onto slides in Reagent 2.

Immunostaining was performed to measure immunoreactive phosphorylated-S129 α-synuclein and identify patched neurons for morphology analysis. Table 1 below details the antibodies used to label tyrosine hydroxylase (dopaminergic cell marker), Aldh1a1 (marker of ventral tier SNc neurons), pS129 α-synuclein, and Neurobiotin-filled cells.

**Table 1.**
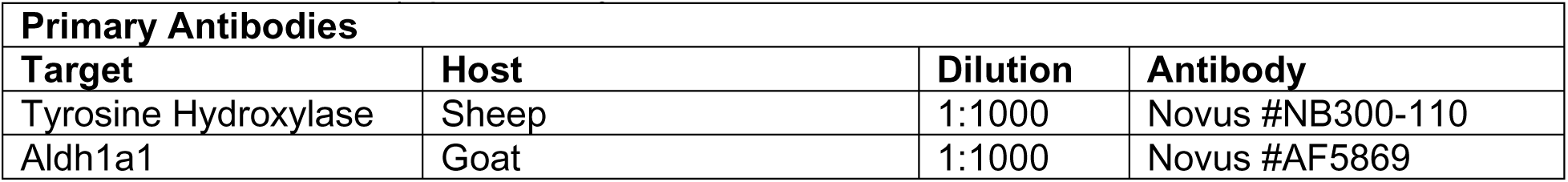
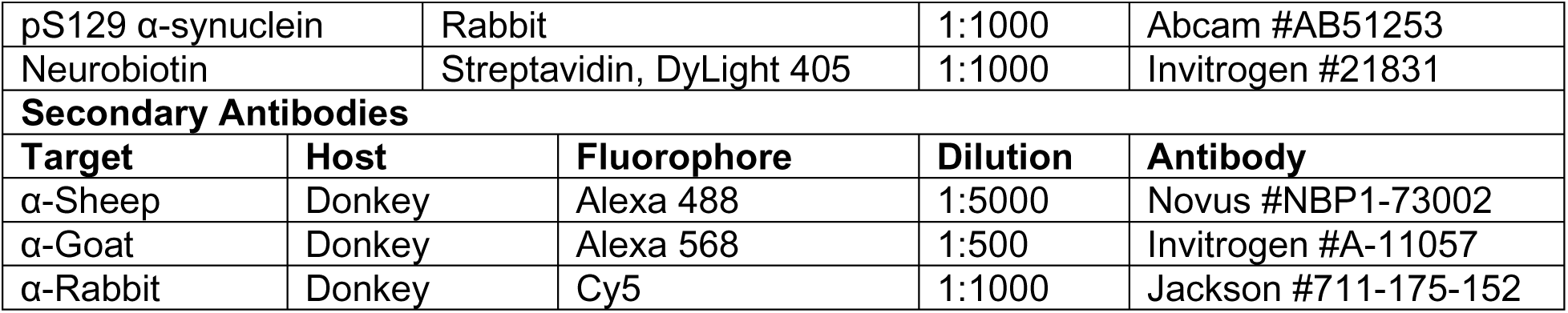
Antibodies for immunohistochemistry.

| Primary Antibodies |  |  |  |
| --- | --- | --- | --- |
| Target | Host | Dilution | Antibody |
| Tyrosine Hydroxylase | Sheep | 1:1000 | Novus #NB300-110 |
| Aldh1a1 | Goat | 1:1000 | Novus #AF5869 |

**Table 1. Antibodies for immunohistochemistry.**
|  |  |  |  |  |
| --- | --- | --- | --- | --- |
| pS129 α-synuclein | Rabbit | 1:1000 | Abcam #AB51253 | |
| Neurobiotin | Streptavidin, DyLight 405 | 1:1000 | Invitrogen #21831 |  |
| Secondary Antibodies |  |  |  |  |
| Target | Host | Fluorophore | Dilution | Antibody |
| α-Sheep | Donkey | Alexa 488 | 1:5000 | Novus #NBP1-73002 |
| α-Goat | Donkey | Alexa 568 | 1:500 | Invitrogen #A-11057 |
| α-Rabbit | Donkey | Cy5 | 1:1000 | Jackson #711-175-152 |

### *In situ* hybridization RNAscope

*In situ* hybridization was performed on 16 µm thick midbrain slices from fresh-frozen mouse brain cut on a cryostat. All RNAscope reagents used are commercially available from Advanced Cell Diagnostics (ACD) and procedures for the RNAscope process were followed as recommended by ACD. Probes used were mouse Cacna1g-channel 1 (ACD #582171), Cacna1h-channel 1 (ACD #582181), Cacna1i-channel 1 (ACD #459781), Aldh1a1-channel 2 (ACD # 491321-C2), Calbindin-channel 2 (ACD # 428431-C2), and TH-channel 3 (ACD #317621-C3). TSA Vivid dyes were used at a dilution of 1:3000, with 650 Cyanine 5.5 assigned to channel 1, 570 Cyanine 3 to channel 2, and 520 Fluorescein to channel 3. All slides were counterstained with DAPI before mounting with ProLong Gold Antifade Mountant (Invitrogen #P36934).

### Confocal imaging

Immunostained slices from electrophysiology and RNAscope experiments were imaged using a Leica SP8AOBS++ confocal microscope in the Microscopy and Imaging Shared Resource core facility at Georgetown University. Slices were imaged as tiled z-stacks to identify locations of patched cells for neuronal reconstructions and to assess levels of α-synuclein phosphorylation. Tiled images of a single z plane were acquired for RNAscope analysis. All images were acquired with a 40x oil-immersion objective lens.

### Neuronal reconstructions

Tiled z-stacks (1 µm step height) of Neurobiotin-filled neurons were acquired with confocal imaging and uploaded into Neurolucida software. The somas and dendrites of filled neurons were reconstructed by a semi-automatic process. If identified, axons were not reconstructed. Data for each cell were exported from the software and imported into Igor (Wavemetrics) figure preparation. [Morphological reconstructions will be made available in Neuromorpho.org upon publication of this manuscript]

### Quantification of immunoreactive pS129 α-synuclein

Tiled z-stacks acquired for neuronal reconstructions were analyzed to quantify α-synuclein inclusions within the SNc. In ImageJ, a stack of 50 images (1 µm step) surrounding the location of neurobiotin filled cells were identified and a z-projection of maximum intensity was created. A manual region of interest (ROI) was drawn to outline the borders of the SNc within the image. The Analyze Particles tool was used with a minimum particle size of 3 pixels to count the number of α-synuclein inclusions within the ROI. Data were analyzed in Igor, where the number of inclusions was normalized to the area of the ROI.

### RNAscope image analysis

Tilescan images from confocal imaging were analyzed in ImageJ using a custom macro. A region of interest was drawn around the SNc based on the location of dopaminergic neurons. Average fluorescence within this user-defined region of interest for each CaV mRNA channel. Data were uploaded into Igor Pro where they were further analyzed using custom made procedures.

### Statistical analyses

Statistical analyses made using GraphPad Prism. Statistical significance in two group comparisons was determined using Wilcoxon rank-sum tests (WRST, unpaired). Statistical significance in three or more group comparisons was determined using Kruskal-Wallis tests followed by Dunn’s multiple comparisons tests. Fisher’s exact test was used to assess statistical significance of differences in proportion of binary outcomes (e.g. presence or absence of depolarization block). Box plots show median, 25^th^ and 75^th^ percentiles (boxes), and 9^th^ and 91^st^ percentiles (whiskers). Descriptive statistics of box plots are reported as median (interquartile range). Graphs represent averages with shading indicating ± standard error of the mean (SEM) and descriptive statistics are reported as such. In the text, n indicates the number of cells and N indicates the number of mice, unless otherwise specified.

## Acknowledgements

This work was supported by the Persimmon Foundation, the Michael J Fox Foundation #MJFF-022863, a Parkinson’s Foundation Stanley Fahn Junior Faculty Award #PF-SF-JFA-1040267, and NINDS #R01NS142876 awarded to RCE; and by a National Institute of General Medical Sciences T32 predoctoral fellowship GM142520 awarded to MLB.

