## Supplemental Figures for "α-Synuclein pathology differentially alters T-type calcium currents in vulnerable and resilient substantia nigra dopaminergic subpopulations"

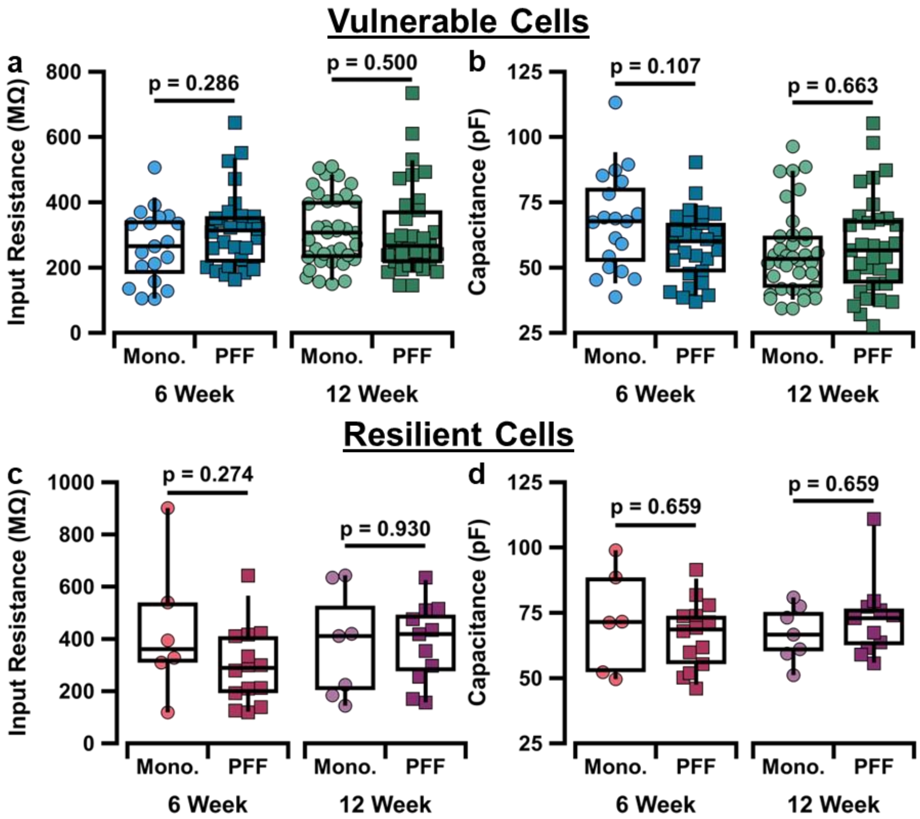


***Supplemental Figure 1. Intrinsic membrane properties of SNc neurons in an α-syn PFF model.*** **a,** Box plots comparing 6-week (left) and 12-week (right) measures of input resistance between monomer and PFF groups in vulnerable cells of the SNc. **b,** Box plots comparing 6-week (left) and 12-week (right) measures of capacitance between monomer and PFF groups in vulnerable cells of the SNc. **c,** Box plots comparing 6-week (left) and 12-week (right) measures of input resistance between monomer and PFF groups in resilient cells of the SNc. **d,** Box plots comparing 6-week (left) and 12-week (right) measures of capacitance between monomer and PFF groups in resilient cells of the SNc.

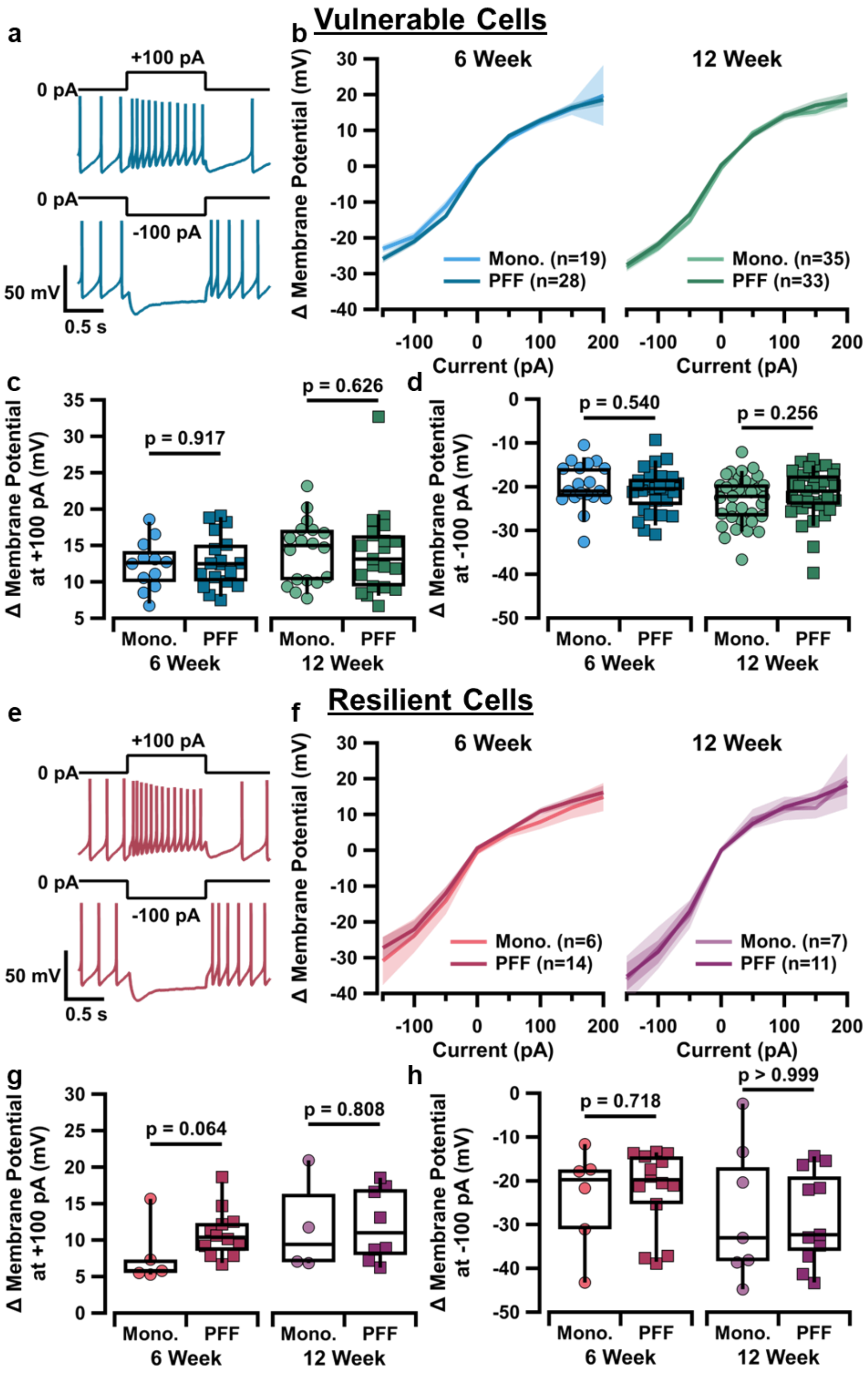


***Supplemental Figure 2. Integration of excitatory and inhibitory inputs by SNc neurons in an α-syn PFF model.*** **a,** Sample traces of current-clamp protocol at +100 pA (top) and -100 pA (bottom) injection steps from vulnerable cell of 6-week PFF-injected mouse. **b,** Graph of current-voltage relationship in ADP cells of monomer and PFF groups at 6 weeks (left) and 12 weeks (right) post-injection. **c,** Box plots comparing change in membrane potential at +100 pA current injection step in vulnerable cells between monomer and PFF groups at 6-weeks (left) and 12-weeks (right) post-injection. **d,** Box plots comparing change in membrane potential at -100 pA current injection step in vulnerable cells between monomer and PFF groups at 6-weeks (left) and 12-weeks (right) post-injection. **e,** Sample traces of current-clamp protocol at +100 pA (top) and -100 pA (bottom) injection steps from resilient cell of 6-week PFF-injected mouse. **f,** Graph of current-voltage relationship in resilient cells of monomer and PFF groups at 6 weeks (left) and 12 weeks (right) post-injection. **g,** Box plots comparing change in membrane potential at +100 pA current injection step in resilient cells between monomer and PFF groups at 6-weeks (left) and 12-weeks (right) post-injection. **h,** Box plots comparing change in membrane potential at -100 pA current injection step in resilient cells between monomer and PFF groups at 6-weeks (left) and 12-weeks (right) post-injection.

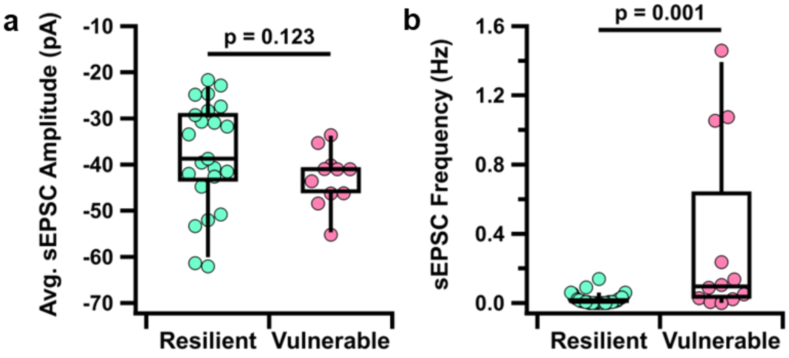


***Supplemental Figure 3. Vulnerable SNc neurons receive more frequent sEPSCs.*** **a,** Box plot comparing average sEPSC amplitude in resilient vs vulnerable cells from monomer-injected animals. [Resilient: -38.7 (16.5) pA, n = 23 vs Vulnerable: -41.0 (6.0) pA, n = 11, p = 0.123, Mann-Whitney test] **b,** Box plot comparing sEPSE frequency in resilient vs vulnerable cells from monomer-injected animals. [Resilient: 0.006 (0.029) Hz, n = 34 vs Vulnerable: 0.096 (0.826) Hz, n = 12, p = 0.001, Mann-Whitney test]
